# Neuromuscular Signals Shape Fatigue and Effort-Based Decision-Making in Humans

**DOI:** 10.64898/2025.12.06.692763

**Authors:** Agostina Casamento-Moran, Aram Kim, Joonhee Leo Lee, Vikram S. Chib

**Affiliations:** Department of Biomedical Engineering, Johns Hopkins School of Medicine; Baltimore, MD, USA; Kennedy Krieger Institute; Baltimore, MD, USA; Kavli Neuroscience Discovery Institute, Johns Hopkins University; Baltimore, MD, USA

## Abstract

Physical fatigue influences our willingness to undertake effortful actions, yet the physiological signals driving this process are not well understood. We created a biofeedback paradigm to distinguish the effects of reduced muscle-force capacity from compensatory increases in neuromuscular activity on effort-based decision-making. Human participants made risky choices about prospective physical effort before and after repeated fatiguing exertions under two biofeedback conditions: force biofeedback, which required increased neuromuscular drive, and EMG biofeedback, which constrained the increase in neuromuscular drive. Both biofeedback conditions led to similar reductions in muscle strength and feelings of fatigue. However, we found that the Force biofeedback condition, which required compensatory neuromuscular activation, produced a significantly greater increase in the subjective cost of effort, thereby altering individuals’ effort-based decision-making more than EMG biofeedback. These findings demonstrate how muscle physiological processes influence feelings of fatigue and decisions to exert effort. Suggesting that fatigue may consist of separate components that collectively motivate behavior while simultaneously protecting and restoring bodily homeostasis during physical challenge.

## Introduction

Fatigue is pervasive in health and disease, significantly impacting our decisions to engage in effortful daily activities^1–3^. For example, if we feel tired after a long, hard day at work, we may decide to skip our evening exercise routine. Previous studies have provided a neurobiological account of how fatigue modulates decisions to engage in effortful physical activity, demonstrating that repeated fatiguing exertions change individuals’ motor cortical state and inflate their subjective valuation of effort (i.e., making effort more costly)^4–7^. These studies primarily focused on understanding brain activity related to the valuation of effort during rest and fatigue and did not investigate how exertion-induced changes in neuromuscular activity contribute to the subjective valuation of effort and feelings of fatigue.

Repeated fatiguing exertions result in peripheral (i.e., muscular^8–12)^, and central (i.e., spinal^13,14^ and supraspinal^13^) changes that lead to, and compensate for, reduced muscle-force capacity. Exertion of effortful contractions trigger a cascade of metabolic and contractile changes within the muscle that limit its ability to generate force^10,15^, a phenomenon known as fatigability^1,2^. To compensate for reduced muscle-force capacity and maintain performance, there is an increase in neuromuscular activity^10,13^. As a result, when fatigued, greater neuromuscular activity is used to generate the same amount of force as when rested. Despite a great deal of work regarding muscle physiology, we have a limited understanding of how the mechanisms underlying muscle-force generation and fatigability influence decisions to exert effort, feelings of fatigue, and subjective assessments of the exerted effort.

It has recently been proposed that fatigue-induced inflations in the subjective valuation of physical effort may reflect a behavioral strategy that promotes rest and recovery in the presence of bodily dyshomeostasis^4,16–18^. In this framework, it is possible that the decrease in muscle-force capacity that occurs with repeated fatiguing exertions may inflate the subjective valuation of effort and trigger feelings of fatigue. On the other hand, increased neuromuscular activity has been proposed to underpin heightened effort perception with repeated fatiguing exertions^19–21^, suggesting that the compensatory increase in neuromuscular activity may inflate the subjective valuation of effort and feelings of fatigue. It is also possible that fatigue-induced changes in muscle force capacity and neuromuscular activity interact to inflate the subjective valuation of effort, give rise to feelings of fatigue, and heighten retrospective assessments of the exerted effort.

## Results

To test these alternative hypotheses, we designed a biofeedback paradigm that separates reductions in muscle force capacity from compensatory increases in neuromuscular activity. We employed a repeated-measures crossover design with two experimental conditions: one where participants received biofeedback directly proportional to the force they exerted (Force biofeedback), and another where biofeedback was proportional to the electromyography (EMG) amplitude associated with their grip exertion (EMG biofeedback). During the Force biofeedback condition, participants repeatedly exert a prescribed amount of force. To compensate for the reduced muscle-force capacity across the session, they must increase neuromuscular activity (**Fig. 1A**). Conversely, in the EMG biofeedback condition, we prevent compensatory increases in neuromuscular activity by having participants maintain a consistent level of EMG activity (**Fig. 1B**). In this setting, the decline in muscle-force capacity results in participants exerting lower forces as the session progresses. This manipulation allowed us to independently evaluate the impacts of decreased muscle-force capacity and increased neuromuscular activity on the subjective valuation of effort (i.e., decision-making regarding prospective effort), fatigue ratings (i.e., feelings of fatigue), and effort assessments (i.e., retrospective assessment of previously exerted force). If compensatory increases in neuromuscular activity mainly influence the subjective valuation of effort, feelings of fatigue, and retrospective effort assessments, then we expect that only the Force biofeedback condition—not the EMG biofeedback—would alter effort valuation, induce feelings of fatigue, and increase effort ratings. Conversely, if the primary factor is the decrease in muscle-force capacity, we predict that changes in effort valuation, feelings of fatigue, and effort assessments would be similar across both biofeedback conditions.

**Figure 1.**
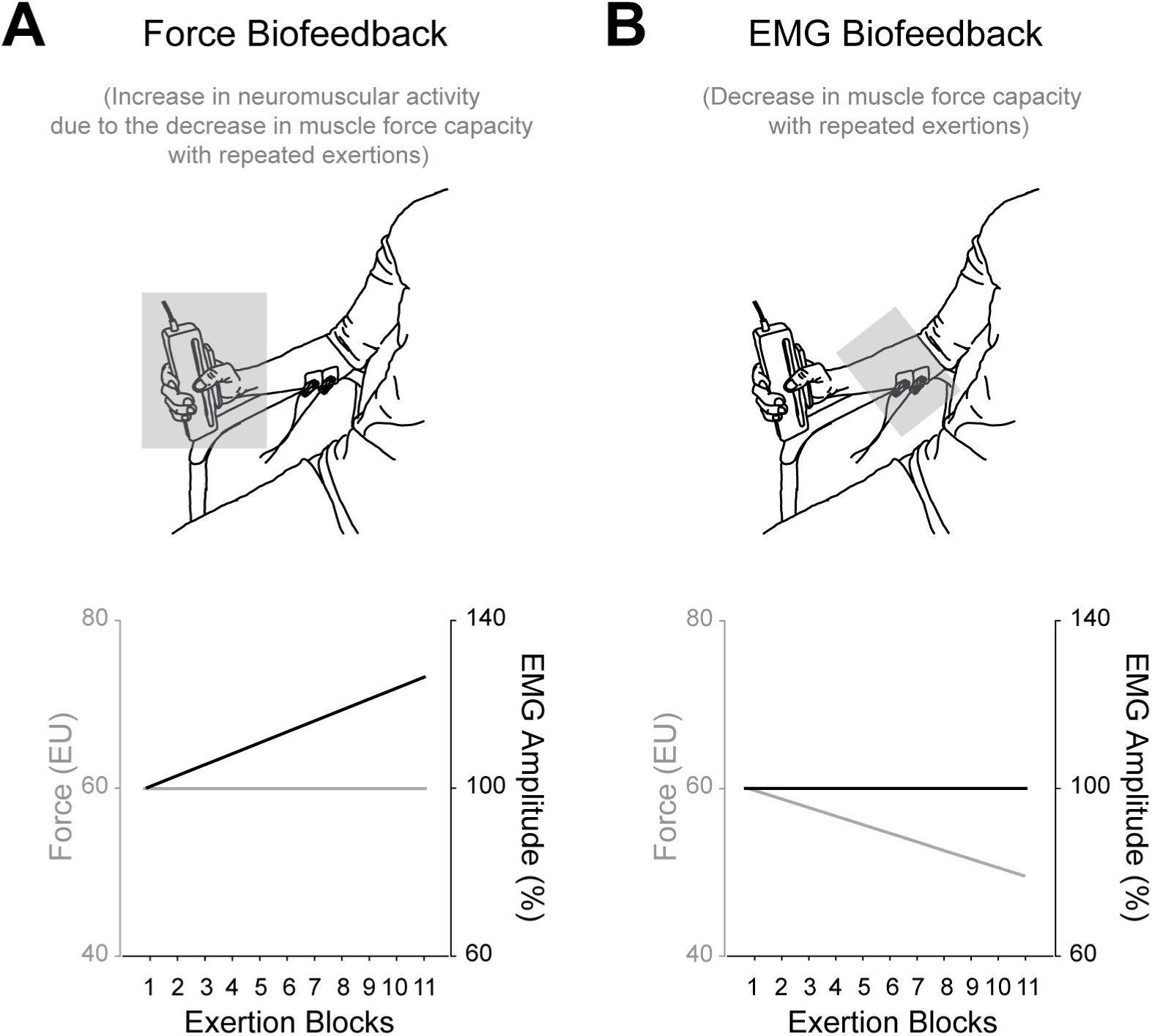
Hypothesized Force and EMG responses during the biofeedback conditions. **A)** During force biofeedback, participants must exert a prescribed amount of force, which increases neuromuscular activity (i.e., EMG amplitude). In this condition, both a decrease in muscle force capacity and a compensatory increase in neuromuscular activity occur. **B)** During EMG biofeedback, participants must maintain a prescribed level of neuromuscular activity, which results in lower exerted force due to the decrease in muscle-force capacity. This condition isolates reduced muscle-force capacity from the compensatory increase in neuromuscular activity.

To investigate how decreases in muscle-force capacity and compensatory increases in neuromuscular activity influence decisions to exert effort, participants made risky choices about prospective effort before and after bouts of effortful exertion. The first session of choices was used to characterize participants’ subjective valuation of effort in a baseline, rested state (Effort-based Choice; **Fig. 2A**). After this *baseline choice phase*, participants performed a *fatigue choice phase* that consisted of alternating between blocks of effort-based choices and blocks of exertion trials (n=10; **Fig. 2A, B**). All the choices were for prospective effort. To ensure that participants’ decisions had actual consequences, 10 trials were randomly selected at the end of the experiment for the participants to play out. In addition, we examined how reduced muscle-force capacity and the compensatory increase in neuromuscular activity influence fatigue ratings and the retrospective assessment of effort. To do so, we asked participants to rate their fatigue after each choice and exertion block (Fatigue Rating; **Fig. 2C**) and the effort they perceived to have just exerted after each exertion block (Effort Assessment; **Fig. 2D**). See **Fig. 2E** for a graphical representation of the experimental schedule.

**Figure 2.**
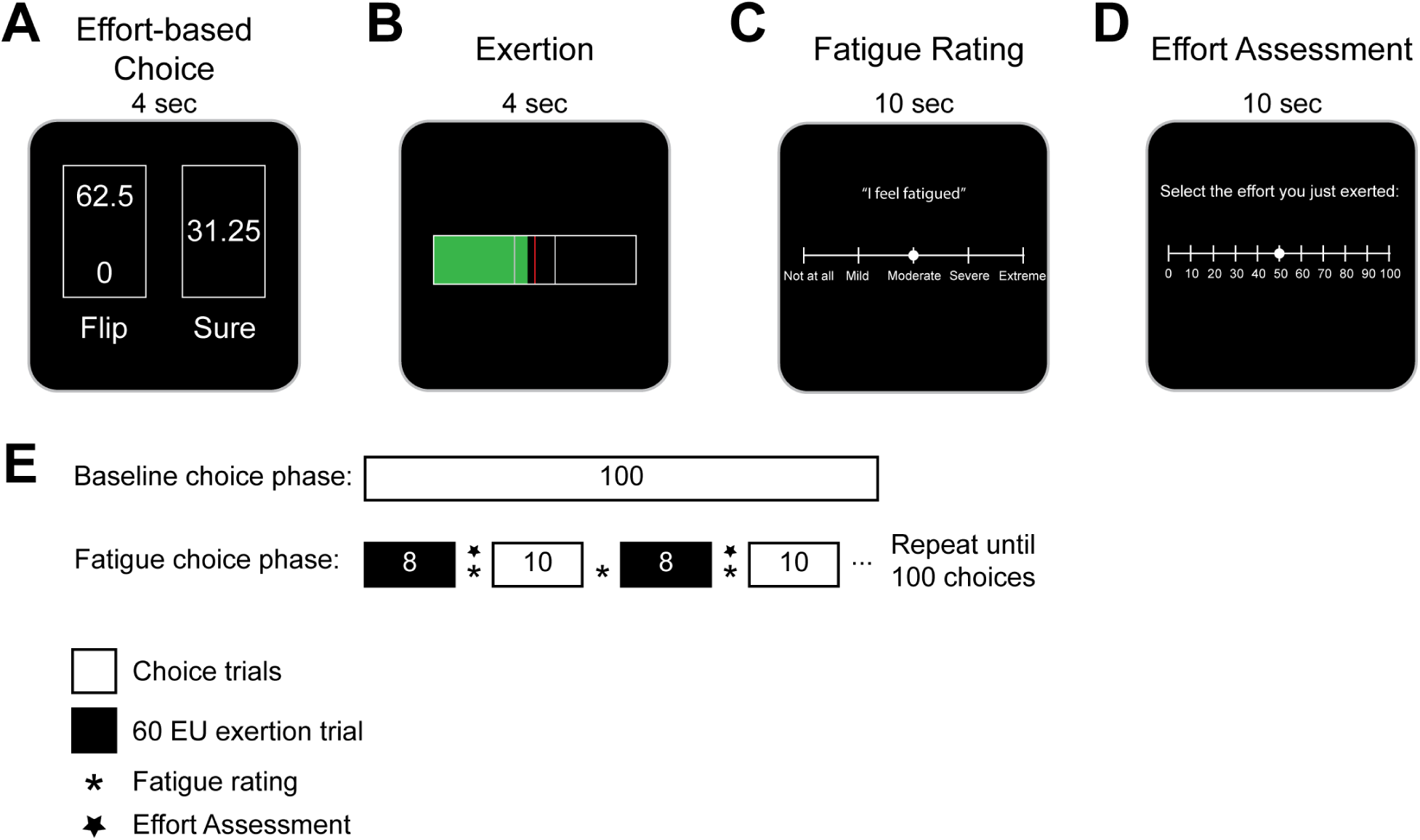
Experimental design. **A)** During effort-based choices participants were presented with a series of risky decisions between 2 options: exerting a low amount of effort with certainty (‘Sure’) or taking a gamble that could result in either a higher level of exertion or no exertion with equal probability (‘Flip’). The effort amounts were presented on a 0 to 100 scale that participants were trained on during an association phase prior to choice. An effort level of zero corresponded to no exertion and 100 to 80% of a participant’s maximum voluntary capacity (MVC). **B)** To investigate how reduced muscle-force capacity and the compensatory increase in neuromuscular activity influence effort-based decision-making, blocks of exertion trials were interspersed with blocks of effort-based choices. Two types of exertion trials (Force and EMG Biofeedback) were presented on separate days. During Force Biofeedback trials, participants exerted grip force on a hand dynamometer, and grip forces from the dynamometer directly related to the filling of a horizontal bar, where the center of the bar corresponded to an effort level of 60. During EMG Biofeedback trials, participants exerted grip force on the hand dynamometer, and EMG signals from exertion filled the horizontal bar, where the center of the bar corresponded to the EMG signals associated with the amount of EMG used when exerting an effort level of 60. **C)** Following exertion blocks, we asked participants to report the level of effort they perceived to have just exerted. **D)** Before and after each exertion block, we also surveyed participants about the level of fatigue that they experienced. **E)** Experimental schedule. The experiment was divided into baseline and fatigue choice phases. The *baseline choice phase* consisted of 100 effort-based choices presented in a randomized order. Following the baseline choice phase, participants performed the *fatigue choice phase* of the experiment, in which they underwent repeated exertion trials (indicated in black) to induce fatigability. On separate days, participants performed either the Force Biofeedback trials or EMG Biofeedback trials during the fatigue phase. While exertion feedback differed on these days, the schedule of choice and exertion trials were identical. The fatigue choice phase began with participants alternating between exertion trials (black) and blocks of choice trials (white). Each exertion block lasted until participants exerted eight successful trials by achieving the target level. Each choice block consisted of 10 effort-based choices randomly sampled from the same set used in the baseline choice phase. At the end of each exertion block, participants reported the level of effort they perceived to have exerted (panel **C**; star) and the level of fatigue that they experienced (panel **D**; asterisk).

Participants completed two experimental sessions, separated by at least 24 hours, that were identical except for the biofeedback provided during exertion blocks in the *fatigue choice phase*. Specifically, during Force and EMG biofeedback conditions, participants had to exert 8 successful trials at either a force or EMG level of 60 Effort Units (EU), respectively (see Methods for details). The order in which participants performed each biofeedback condition was randomly assigned. Participants were informed that both experimental sessions were identical. Importantly, participants did not perceive the difference between the two biofeedback conditions, as assessed with questionnaires at the end of the second session (see Methods for details).

### Force and EMG biofeedback differentially influence muscle force capacity and compensatory increases in neuromuscular activity

To evaluate the effectiveness of our experimental manipulation in distinguishing between the decrease in muscle force capacity and the compensatory increase in neuromuscular activity, we assessed the magnitude of exerted force and EMG during each biofeedback condition. During the Force Biofeedback condition, participants accurately performed repeated exertions at 60 EU, and their EMG amplitude increased from the first to the last exertion block (One-sample t-test: *t_(23)_ =* 2.99, *p* = 0.006; **Fig. 3A, B**). In contrast, during the EMG Biofeedback condition, participants maintained the EMG amplitude on target across repeated exertions, while the force they produced decreased from the first to the last exertion block (One-sample t-test: *t_(23)_ =* -3.57, *p* = 0.001; **Fig. 3C, D**). These findings demonstrate that the Force and EMG biofeedback conditions successfully induced distinct compensatory responses in neuromuscular activity and muscle-force capacity. To determine whether the biofeedback conditions reduced muscle-force capacity to a similar degree (i.e., caused comparable levels of fatigability), we measured participants’ maximal voluntary force capacity (MVC) before and after the *fatigue choice phase* during both sessions. We observed that while both biofeedback conditions reduced MVC (Linear mixed-effects model: *t_(92)_ =* -1.98, *p =* 0.049), the reduction was similar across conditions, as indicated by a non-significant biofeedback by time point interaction (Linear mixed-effects model: *t_(92)_ =* -0.54, *p =* 0.58, see Methods for details). These results suggest that the EMG biofeedback condition prevented the compensatory increase in neuromuscular activity while inducing a comparable reduction in muscle-force capacity as the Force biofeedback condition.

**Figure 3.**
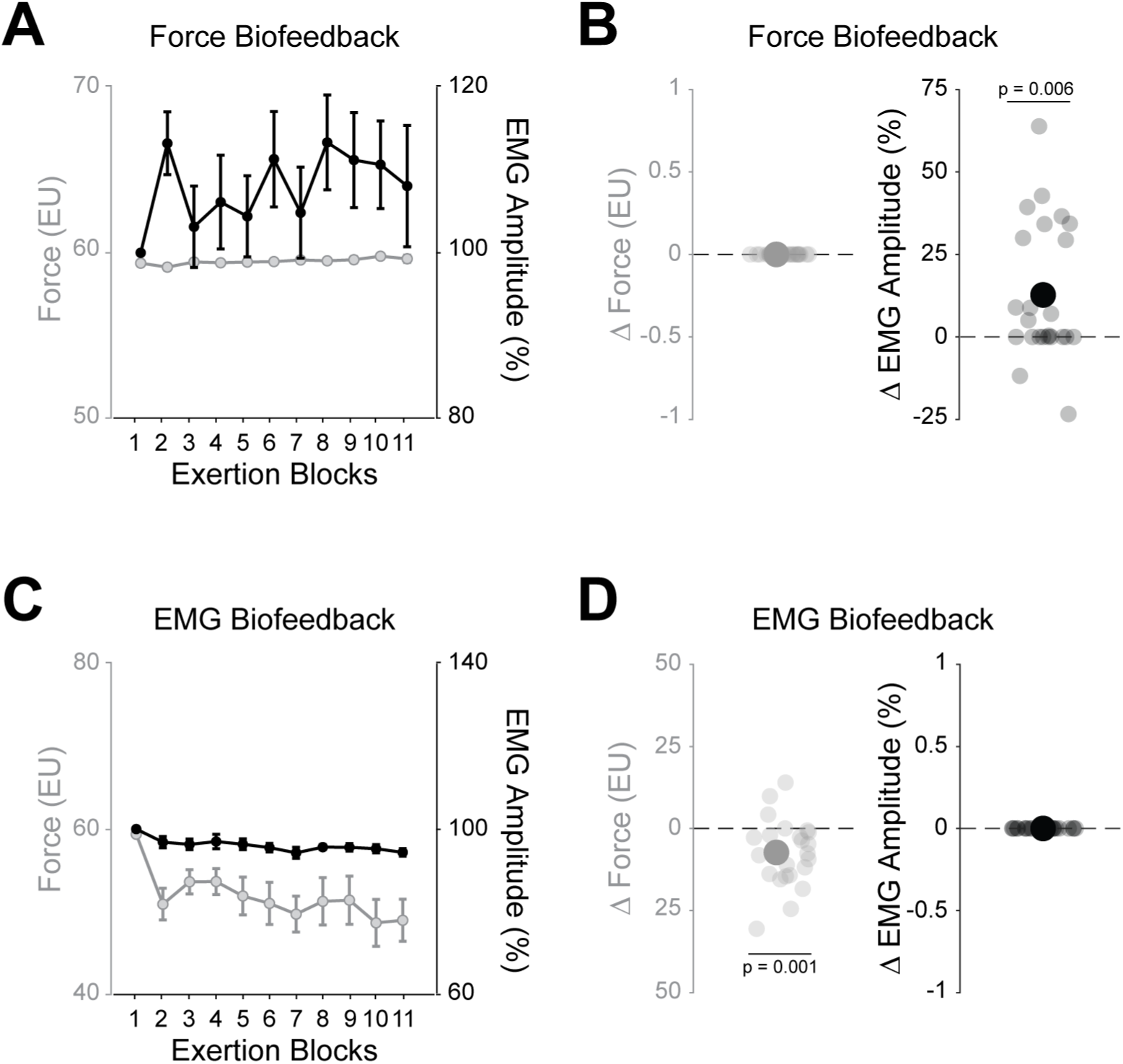
Force and EMG responses during the biofeedback conditions. **A)** Illustration of the group-mean exerted force and EMG amplitude as a function of exertion block in the Force Biofeedback condition. Repeated exertion blocks showed participants increasing their EMG amplitude to maintain the target force. Error bars indicate SEM. **B)** Difference in exerted force (left panel) and EMG amplitude (right panel) between the first and last exertion blocks of the Force Biofeedback condition. Although participants accurately maintained the target force, their EMG amplitude increased significantly. The large point indicates the group average, while small points reflect individual participant means. **C)** Illustration of the group-mean exerted force and EMG amplitude as a function of exertion block in the EMG Biofeedback condition. Through repeated exertion blocks, participants maintain their EMG amplitude while their exerted force decreases. Error bars indicate SEM. **D)** Difference in exerted force (left panel) and EMG amplitude (right panel) between the first and last exertion blocks of the EMG Biofeedback condition. While participants accurately maintained the target EMG, their exerted force decreased significantly. The large point indicates the group average, and small points reflect individual participant means.

### Force and EMG biofeedback differentially increase the subjective value of effort

We used participants’ effort choice data to characterize their subjective value of effort, employing a subjective cost function 𝑉𝑉(𝑥𝑥) = −(−𝑥𝑥)^𝜌𝜌^, where 𝑥𝑥 ≤ 0 and 𝑉𝑉 represents the subjective cost of an objective effort level 𝑥𝑥. ρ is a participant-specific parameter that describes how an individual subjectively represents the effort level 𝑥𝑥. In this model, 𝜌𝜌 is flexible enough to capture increasing, decreasing, or constant marginal changes in subjective effort valuation as absolute effort levels increase. When 𝜌𝜌 = 1, it indicates that a participant’s subjective effort cost matches the absolute effort levels, with no bias toward choosing risky or sure options when the prospects have equal expected value. A 𝜌𝜌 less than 1 suggests decreased sensitivity to changes in subjective effort cost as effort increases, biasing the participant toward selecting the risky effort option when the prospects have equal expected values. Conversely, a 𝜌𝜌 greater than 1 indicates greater sensitivity to changes in effort, biasing the participant toward choosing the sure effort option when the prospects have equal expected values. We applied this model to participants’ effort choices to examine how the subjective value of effort is represented in a rested state and after fatiguing exertions^4,22^.

During the *fatigue choice phase* in both biofeedback conditions, we observed that repeated fatiguing exertions led to greater risk aversion toward effort. This effect was more pronounced with Force biofeedback than EMG biofeedback. Specifically, we used a repeated measures ANOVA to analyze how subjective effort cost changed from the baseline (ρ_Baseline_) to the fatigue choice phase (ρ_Fatigue_) and across biofeedback conditions. We found a significant main effect of fatigue (repeated-measures ANOVA: *F_(1,23)_ =* 15.03, *p <* 0.001), indicating that ρ_Fatigue_ was significantly higher than ρ_Baseline_ across both biofeedback conditions. Additionally, there was a significant interaction between biofeedback conditions and fatigue (repeated-measures ANOVA: *F_(1,23)_ =* 6.41, *p =* 0.019; **Fig. 4A, B**), showing that Force biofeedback caused a greater exertion-induced increase in ρ than EMG biofeedback.

**Figure 4.**
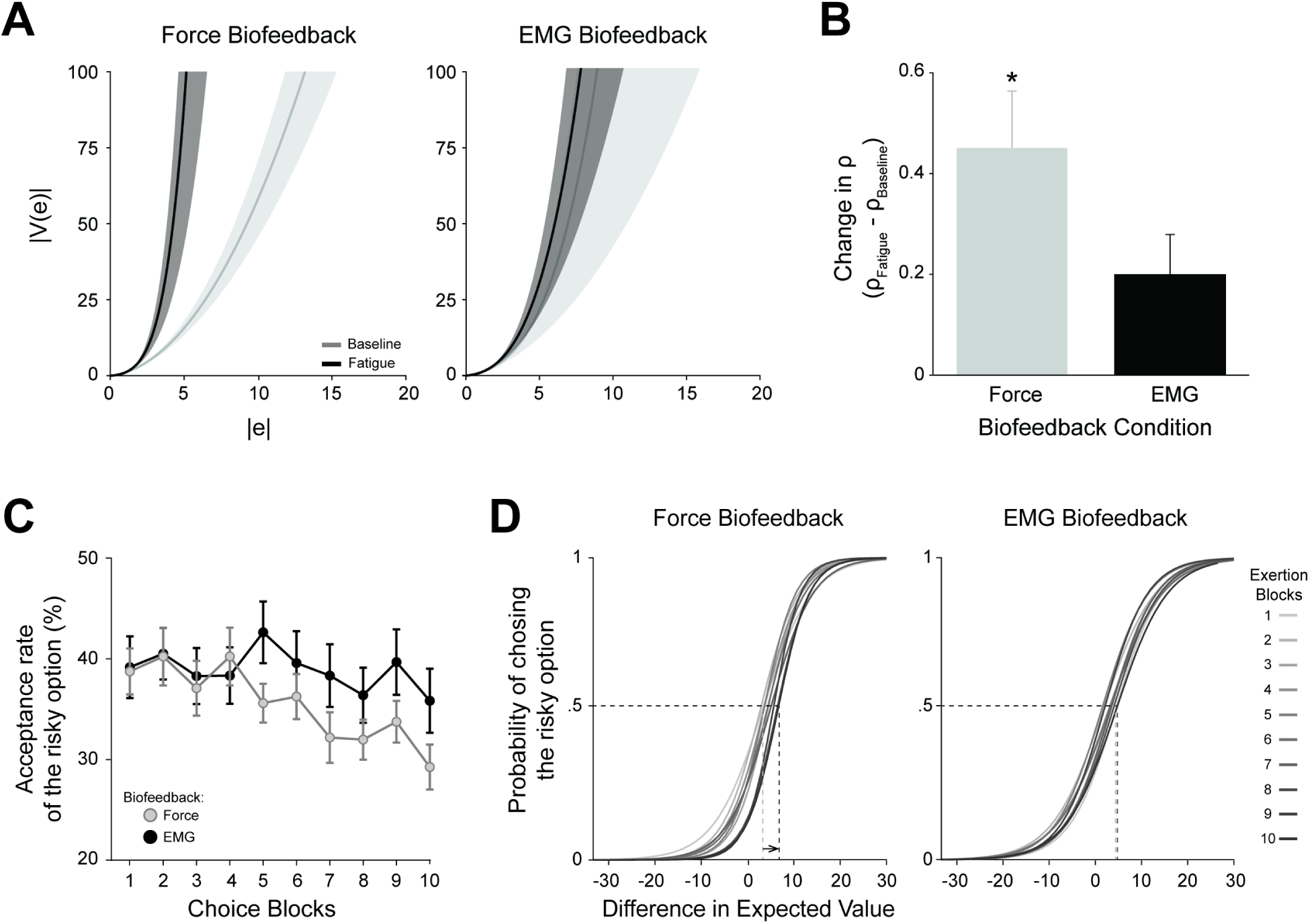
Effort-based decision-making data. **A)** The function used to model the subjective cost of effort is expressed as V(x) = −(−x)^ρ^. Effort cost functions during Force (left) and EMG (right) Biofeedback conditions, using mean values of the ρ estimates, are shown with solid lines (baseline: light gray; fatigue: dark gray), with SEM indicated by the shaded regions. Repeating effortful exertions increases the marginal cost of effort more during Force biofeedback than during EMG biofeedback. To better illustrate the cost functions, the x- and y-axes shown are not on the same scale. **B)** The effort subjectivity parameter (ρ) increased to a greater extent between the baseline and fatigue choice phases in the Force biofeedback compared to EMG biofeedback conditions. Error bars indicate SEM. Paired t-test (two-tailed): *p = 0.019. **C)** Participants’ acceptance rate of the risky option across choice blocks during the Force (gray) and EMG (black) biofeedback conditions. Acceptance rates remained stable over time with EMG biofeedback but decreased gradually with Force biofeedback. Error bars indicate SEM. **D)** Probability of choosing the risky option as a function of the difference in expected value between the risky and sure, shown separately for Force and EMG biofeedback conditions. For visualization, each psychometric curve (in a gradient of gray) represents the average curve for all participants within each choice block (i.e., 10 choices per block). Compared to the EMG condition, Force biofeedback caused more pronounced rightward shift (black arrow) in the psychometric curves over the course of exertion blocks, indicating that individuals became more risk averse for effort as exertions blocks increased.

We also conducted a model-free analysis of changes in choice behavior (i.e., the proportion of accepted risky options) which supported the finding of a more significant decrease in risk aversion for effort in the Force compared to the EMG biofeedback condition. We evaluated how the acceptance rate of the risky option varied over time as a function of biofeedback condition, the number of choice blocks, and their interaction. The acceptance rate of the risky option decreased more in the Force biofeedback condition compared to the EMG biofeedback condition (Linear mixed-effects model: *t_(434)_ =* 2.06, *p =* 0.04; **Fig. 4C**). Additionally, we analyzed participants’ choices based on biofeedback, trial number, and differences in expected value, including their interactions (see Methods for details). Our analysis revealed that the Force biofeedback condition produced a significantly larger and more progressive shift in effort preferences across blocks compared to the EMG biofeedback condition (Mixed-effect logistic regression: *t_(4361)_ =* 3.18, *p =* 0.001; **Fig. 4D**). The fact that both biofeedback conditions increased the subjective valuation of effort and that this effect was most pronounced with Force biofeedback indicates that the reduction in muscle-force capacity, combined with a compensatory rise in neuromuscular activity, collectively influences individuals’ effort-based decisions.

Because the biofeedback conditions occurred on different days, we also confirmed that there were no differences in the subjective value of effort at rest across days. Specifically, we checked whether participants’ subjective valuation of effort during the *baseline choice phase* was consistent across biofeedback conditions (i.e., before participants experienced biofeedback-induced fatigue). We found that participants showed similar choice behavior in the baseline phase (ρ_Baseline_) across both biofeedback conditions, as indicated by similar ρ_Baseline_ parameters (two-tailed paired t-test: *t_(23)_ =* 0.38, *p =* 0.71; **Supplementary Fig. 1**). This finding suggests that differences in the subjective valuation of effort were caused by the biofeedback conditions themselves, rather than changes in effort valuation at rest across days.

### Fatigue ratings increased similarly for both biofeedback conditions

We next examined how reduced muscle-force capacity and the compensatory increase in neuromuscular activity affect feelings of fatigue by modeling participants’ Fatigue Ratings during the *fatigue choice phase* as a function of biofeedback condition, block number, and their interaction. We found that while fatigue ratings increased across blocks (Linear mixed-effects model: *t_(1004)_ =* 11.02, *p <* 0.001), there was no significant difference between biofeedback conditions (Linear mixed-effects model: *t_(1004)_ = -*1.53, *p =* 0.13). Additionally, the rate at which fatigue ratings increased was similar between the two biofeedback conditions (Linear mixed-effects model: *t_(1004)_ =* 0.64, *p =* 0.52; **Fig. 5A**). These findings indicate that the decline in muscle-force capacity is likely the main factor driving fatigue ratings, supporting the idea that subjective feelings of fatigue reflect an affective response to internal physiological imbalance^16,18,23^. Furthermore, they suggest at least a partial dissociation between subjective fatigue sensations and fatigue-related changes in effort-based decision-making.

**Figure 5.**
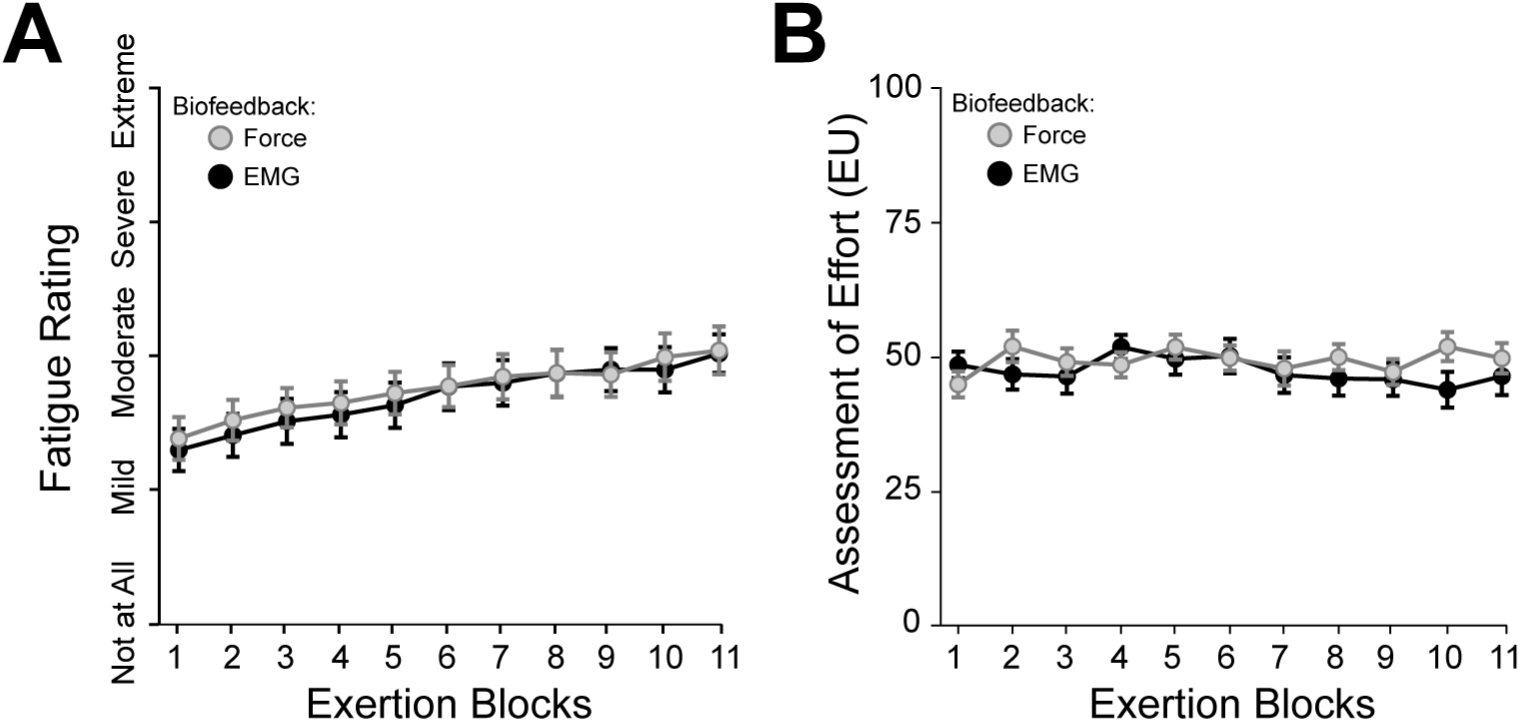
Fatigue Ratings and Effort Assessments. **A)** Participants’ fatigue ratings as a function of the exertion block. Participants’ fatigue ratings increased similarly across exertion blocks in both the Force and EMG Biofeedback conditions. Error bars indicate SEM. **B)** Participants’ assessments of effort as a function of the exertion block. Participants’ effort assessments remained relatively constant across exertion blocks in both biofeedback conditions. Error bars indicate SEM.

### Assessments of effort remained unchanged in both biofeedback conditions

Studies that previously identified an association between neuromuscular activity and effort perception led us to test whether effort perception reflects the magnitude of the neural drive to activate the muscles in our experiment^19–21^. Based on prior work, we hypothesized that the compensatory increase in neuromuscular activity with Force biofeedback would elevate effort assessments. Additionally, we hypothesized that preventing the compensatory increase in neuromuscular activity with EMG biofeedback would stop the rise in effort assessment. We modeled participants’ Effort Assessments during the *fatigue choice phase* as a function of biofeedback condition, block number, and their interaction. Contrary to our hypotheses, effort assessments remained unchanged across blocks (Linear mixed-effects model: *t_(545)_ =* 0.67, *p =* 0.50) and were similar between biofeedback conditions (Linear mixed-effects model: *t_(545)_ =* 0.52, *p =* 0.60; **Fig. 5B**). To further examine changes in effort assessments, we compared responses before and after the fatigue choice phase across effort levels, using participant responses from assessments performed both before and after (see Supplementary Materials for details). We tested two models (i.e., one for each biofeedback condition) to evaluate effort assessment as a function of target effort level, time point (before and after the *fatigue choice phase*), and their interaction. Once again, effort assessments did not increase across a range of forces (10 to 80 EU) after the *fatigue choice phase* (Linear mixed-effects model: *F_(16,2096)_ =* 0.19, *p =* 0.66; **Supplementary Fig. 2**). These findings suggest that, in this paradigm, neither the decrease in muscle-force capacity nor the compensatory increase in neuromuscular activity affected retrospective effort assessments, indicating at least partially separate processes related to how fatiguing physical exertion influences decisions about prospective effort and retrospective effort evaluation.

## Discussion

This study aimed to differentiate the effects of exertion-induced reductions in muscle-force capacity and compensatory increases in neuromuscular activity on the subjective valuation of effort, feelings of fatigue, and assessments of exerted effort. Using an innovative biofeedback paradigm, we manipulated the relationship between force exertion and neuromuscular activity. Our results indicate that while both Force and EMG biofeedback conditions caused similar increases in feelings of fatigue and no change in effort assessment, the condition requiring compensatory increases in neuromuscular activity (Force biofeedback) significantly elevated the subjective cost of effort compared to the condition where only the decrease in muscle-force capacity occurred (EMG biofeedback). Overall, these findings demonstrate a connection between the physiological factors driving fatigability and those affecting effort-based decision-making, emphasizing that subjective effort costs are more responsive to compensatory neuromuscular activity.

Our results support and refine the dyshomeostasis model of fatigue, which suggests that fatigue is a behavioral response to internal physiological imbalance^16–18^. We observed that feelings of fatigue increased similarly across biofeedback conditions, despite having different neuromuscular profiles and choice behaviors. These results imply that it is the decline in muscle-force capacity (i.e., the internal physiological imbalance), rather than compensatory neural drive, that leads to feelings of fatigue. Conversely, we found that both the decline in muscle-force capacity and compensatory neural drive influence decision-making about effort exertion. These findings align with allostatic models^24–26^, which view fatigue and decisions to exert effort as a regulatory process that promotes rest and recovery to restore bodily homeostasis. In this framework, interoceptive signals (i.e., those related to the decrease in muscle-force capacity) may be implicitly integrated with sensorimotor information (i.e., those related to the increase in neuromuscular drive) to influence decisions about effortful engagement^25,27–29^.

Although both reflect bodily challenges, fatigue ratings and effort-based decisions may be distinct but interacting components that depend on partially separable neurophysiological signals. This may help explain why people sometimes override feelings of fatigue and persist, while in other cases, they opt to rest. Effective effort-based decisions likely depend on an integrated estimate of internal state^25,27–29^, and our findings suggest that neuromuscular information may help interpret imprecise interoceptive cues^30,31^ to influence behavior.

Surprisingly, throughout the experiment, participants’ assessments of effort (i.e., retrospective judgments of exertion) remained stable. This contradicts previous research, which has suggested that for similar levels of exertion, assessments of effort increase when in a fatigued state^32,33^. It is possible that we did not observe a fatigue-induced shift in assessments of effort because we used a submaximal fatiguing task (isometric grip), which may not produce the same perceptual recalibrations seen in previous studies that employed whole-body exercise paradigms^32^. Additionally, our paradigm explicitly separated ratings of fatigue from assessments of exerted effort, whereas earlier work often combined these two measures^32,33^. Our use of well-calibrated effort levels during the association phase may have anchored participants’ expectations and provided a consistent reference for judging exertion. Our effort assessment method contrasts with effort rating scales, such as Ratings of Perceived Exertion (RPE), which rely heavily on individual experience and cardiovascular feedback^33^. Going forward, it will be important to investigate whether retrospective effort assessments are more sensitive to different types or intensities of exertion and how task context influences these judgments.

Understanding the different components of fatigue is especially important in clinical settings, where fatigue is among the most debilitating and least understood symptoms across many disorders^2,3^, including multiple sclerosis^34–36^, stroke^37^, cancer^38,39^, depression^16,40^, and Long COVID^41–43^. Currently, clinical assessments depend on composite fatigue scores that merge various physiological and psychological aspects and mask individual differences in symptom presentation. Our findings indicate that breaking down fatigue into separate components could reveal its underlying mechanisms, enhance diagnostic accuracy, and help develop targeted treatments.

This perspective may also explain why current fatigue interventions (e.g., cognitive behavioral therapy, graded exercise, pharmacological agents) have highly variable efficacy^44–50^. If different clinical populations show different dominant features of fatigue, a one-size-fits-all treatment approach is unlikely to be effective. Instead, a more personalized strategy that targets specific aspects of fatigue might offer greater therapeutic benefits. By combining neuromuscular manipulations with computational modeling, our study shows how different physiological processes influence fatigue-related behavior. These findings emphasize that fatigue may consist of separate components that together serve a regulatory role: encouraging behavior to protect and restore bodily homeostasis in the face of physical challenge.

## Methods

### Experimental setup

Presentation of visual stimuli and data collection were conducted using custom MATLAB (http://www.mathworks.com) scripts implementing the PsychToolBox libraries^51^. A hand clench dynamometer (TSD121B-MRI, BIOPAC Systems, Inc., Goleta, CA) was used to measure grip force. Force signals were amplified 1000 times (DA100C, BIOPAC Systems, Inc., Goleta, CA) and sampled at 1,000Hz (NI-DAQ card, Model PCIe-6321, National Instruments, Austin, TX). Additionally, electromyographic (EMG) activity from the flexor carpi ulnaris (agonist) and extensor carpi radialis (antagonist) was recorded using silver/silver chloride (Ag/AgCl) electrodes (B01AME7YC0 3M 2560) and 30 cm leads (LEAD108C, BIOPAC Systems, Inc., Goleta, CA). EMG signals were band-pass filtered from 1-500 Hz, amplified 1000 times (EMG100C-MRI, BIOPAC Systems, Inc., Goleta, CA), and sampled at 1,000Hz (NI-DAQ card, Model PCIe-6321, National Instruments, Austin, TX).

### Experimental design

#### Participants

Twenty-seven healthy individuals participated in the experiment, three of whom were ultimately excluded from the final analyses because their subjectivity parameter ρ, obtained from the baseline choice phase, was more than two standard deviations from the population mean for that phase. The final analysis included 24 participants in total (mean age, 23 years; age range, 18–36 years; 12 females). All participants were prescreened to exclude those with a prior history of neurological or psychiatric illness. The Johns Hopkins School of Medicine Institutional Review Board approved this study, and all participants provided written informed consent.

#### Experimental paradigm

We employed a repeated-measures crossover design. Participants completed the experimental session twice: once with force biofeedback and once with EMG biofeedback during the *fatigue choice phase*. The order of the biofeedback sessions was randomized among participants, with at least 24 hours between sessions. Participants were unaware of the specific experimental manipulation and were told that both sessions were identical. Importantly, they did not perceive a difference between the biofeedback conditions, as confirmed by questionnaires administered at the end of the second session. Before the first session, participants were informed they would receive a fixed show-up fee of $80 after completing the second session. It was emphasized that this fee was independent of their performance or behavior during the experiment.

The association, assessment, and choice phases of the experiment described below are identical to those we have previously used^4,22^. The experiment started by determining participants’ maximum voluntary capacity (MVC) through selecting the highest force achieved over three consecutive repetitions on the hand-clench dynamometer (MVC_Baseline_). During these repetitions, participants were unaware of the upcoming experimental phases and were instructed to squeeze with their maximum effort. Next, participants completed an *association phase* where they learned to link effort levels (defined relative to MVC) with the force exerted against the dynamometer. Effort units (EU) ranged from 0 (no exertion) to 100 (force equal to 80% of a participant’s MVC). Each training block involved five trials for each target level, varying from 10 to 80 in increments of 10 EU, with training blocks presented in an interleaved order (80, 10, 70, 20, 60, 30, 50, 40) to reduce fatigue and tiredness. To further prevent fatigue during this phase, association trials at the highest effort level (100% MVC) were avoided. Each trial of a training block began with a 1-second display of the target effort level, followed by an effort task with visual feedback in the form of a vertical black bar, akin to a thermometer, which turned white as participants squeezed harder (4 s). The bar’s bottom and top represented effort levels 0 and 100, respectively. Participants were instructed to reach the target zone (± 5 effort units of the target) as quickly as possible and sustain their force within this zone for the entire 4 seconds. The target zone was highlighted in green if the effort was within the specified range, and red otherwise. If participants remained within the target zone for more than two-thirds of the total time (2.67 s) during squeezing, the trial was deemed successful. These success criteria aimed to ensure consistent effort duration across all effort conditions. To minimize fatigue, a fixation cross (2 s) and a ‘Get Ready’ screen (1 s) separated trials within a training block, and participants received 30 seconds of rest after every four training blocks.

Following the *association phase*, participants underwent an *assessment phase* to determine whether they had successfully learned to associate effort levels with the actual force exerted. They were tested on each of the previously trained effort levels (10–80, in increments of 10 EU), six times per level, in a randomized order. Each assessment trial involved displaying a black horizontal bar that participants were instructed to fill by gripping the hand dynamometer—turning the effort feedback from red to green once they reached the target effort level. Unlike in the previous phase, the full bar in this phase did not represent an effort level of 100 but instead indicated the specific target effort level for that trial. Participants were instructed to reach the target zone as quickly as possible, maintain their effort within the target for as long as possible, and get a sense of the effort level they were gripping during exertion (4 seconds). After exerting effort, they were shown a number line ranging from 0 to 100 and asked to select the effort level they believed they had exerted. They did this by pressing the ‘LEFT’ and ‘RIGHT’ arrow keys to navigate to their chosen level, then pressing the “SPACE BAR” to confirm their response. No feedback was provided regarding the accuracy of their selection. Following this *assessment phase, p*articipants performed a single MVC trial (MVC_Post-1_) to evaluate any potential decreases in exertion performance that might have developed after the *association* and *assessment tasks*.

To determine participants’ baseline subjective effort cost, they performed an effort-based decision-making task (i.e., *baseline choice phase*) where they made choices about future effort. Before seeing the effort decisions, participants were told that 10 of their decisions would be randomly selected and played out at the end of the experiment, and that they would need to stay in the experimental room until they successfully completed the chosen exertions (*Outcome task*). Since trials were randomly selected at the end, participants were instructed to treat each effort decision independently. They were shown a series of effort choices between two options on the screen within a 4-second time limit: one option involved exerting a small force (S) for certain (the “sure” option), while the other involved taking a risk, which could lead to either a large exertion (F) or no exertion, each with equal probability (the “flip” option) (**Fig. 2A**). Effort levels were displayed on a 0–100 scale, which participants had experienced during the *association task*. Details of these effort choices can be found in our previous study that used this choice set to model the subjective value of effort in a rested state^22^. Participants made their selections by pressing either the “N” (flip) or “M” (sure) keys on the keyboard with their non-dominant hand. The outcome of the gambles was not revealed after the choice, and participants did not perform the exertion task during this phase. One hundred (100) effort choices were presented in a randomized order. Participants were encouraged to make a choice on every trial, although there was no penalty for failing to choose within the four-second window. Failure to make a choice in time was recorded as a missed trial and was not repeated.

Following the *baseline choice phase*, participants completed a *fatigue choice phase* (**Fig. 2E**), where we manipulated the biofeedback provided. Participants alternated between blocks of effort exertion (**Fig. 2B**) and effort-based choices (**Fig. 2A, E**). Specifically, they first completed two consecutive effort exertion blocks, then alternated between effort-based choice blocks and effort exertion until all 100 choices were made. During the effort exertion blocks, biofeedback was manipulated. Both biofeedback conditions involved exerting 8 successful effort trials with a visual display of a black horizontal bar, centered for 4 seconds (**Fig. 2B**). Participants were instructed to fill the bar by gripping the hand-clench dynamometer to turn the bar from red to green and to maintain the effort within the target (green bar) as long as possible. A trial was successful if the participant maintained the exerted force or EMG on target for more than 50% of the trial’s duration (4 seconds); unsuccessful trials were repeated. In the **Force biofeedback condition**, the bar was controlled by the amount of hand-grip force exerted, with a target effort level of 60 ± 5 EU. In the **EMG biofeedback condition,** the bar was controlled by the EMG generated in the Flexor Carpi Ulnaris muscle. The target EMG level was determined during the first effort exertion block of the *fatigue choice phase*, using the 8 successful trials to establish the mean exerted EMG ± 99% confidence interval, approximately ± 5 EU, for subsequent blocks with EMG biofeedback. The raw EMG signals were rectified and smoothed twice with MATLAB’s *‘smoothdata’* function, using a 300 ms running average. Each choice block comprised 10 trials (**Fig. 2A**). Participants were encouraged to make a choice on each trial, though there was no penalty for failing to respond within four seconds. Participants were informed beforehand that fatigue was likely but asked to exert their best effort on each trial. They were also told that performance during exertion trials was independent of choices and outcomes. After each effort exertion block, participants rated their perceived effort on a number line from 0 to 100 (Fig. 2C), indicating how much effort they believed they exerted by using the ‘LEFT’ and ‘RIGHT’ arrow keys to move a cursor and pressing the “SPACE BAR” to confirm. Prior to each effort or choice block, participants completed a computerized fatigue rating scale asking, “I feel fatigued” (**Fig. 2D**), responding by displacing a cursor horizontally along a scale from 0 to 100, where options ranged from “Not at all” (0) to “Extreme” (100), using the arrow keys and confirming with the “SPACE BAR”. After the *fatigue choice phase*, participants repeated the *MVC* (MVC_Post-2_) and *assessment tasks*. The session concluded with the *Outcome task,* where 10 randomly selected choices required exertion. Participants stayed in the testing area until they completed the exertions for all selected trials.

Participants performed these effort-based tasks while sitting comfortably upright and facing a 20-inch square monitor placed 0.5 meters away at eye level. The participant’s dominant shoulder joint was abducted at approximately 25°, the elbow was flexed around 90°, and the wrist remained in a neutral position (**Fig. 1**).

#### Post-experiment debriefing

At the end of the second experimental session, we asked participants whether they noticed any differences between the sessions. None mentioned realizing that their muscle activity was used to control the biofeedback displayed on the screen. Of the 24 participants whose data were analyzed, 9 reported that during one of the sessions (unknowingly to them, the EMG biofeedback session), the therm-like bar was somewhat more variable and harder to control, but it did not impact their performance. None of the participants reported using a different strategy across days.

### Data Analysis

#### Exerted Force and EMG Analysis

We examined the force and EMG exerted during the *fatigue choice phase* by quantifying the mean exerted force or EMG during the stable middle 1 second of the trial (i.e., 2 to 3 s). We excluded data from the first 2 seconds and the last 1 second of the trial to remove variability caused by (a) delayed initiation; (b) increased effort to reach the target; (c) adjustments made to stabilize on the target; (d) decreased effort at the end of the trial. Exerted force is expressed in effort units (EU), which are calculated relative to each participant’s strength (i.e., 100 EU = 80% MVC). Exerted EMG, meanwhile, is presented relative to the EMG during the first block of the fatigability task, where 100% represents the mean EMG used to exert 8 successful trials at 60 EU, and ±8% represents the percentage equivalent to ±5 EU.

We assessed the effectiveness of the biofeedback manipulation by measuring participants’ ability to maintain exerted force and EMG amplitude within the target range at the start (block 1) and end (block 11) of the fatigue phase. We first calculated the average error from the target (60 EU for force and 100% for EMG). To account for the allowed tolerance during biofeedback (±5 EU for force, ±8% for EMG), we adjusted the observed error using the following piecewise functions.

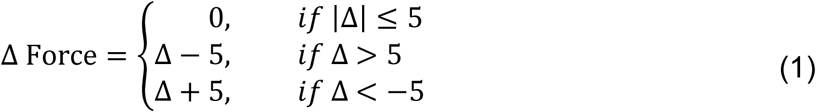

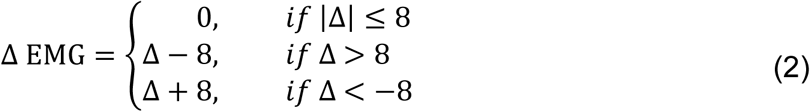

We then examined whether exerted force and EMG differed from target (i.e., 0 after adjustment) using a one-sample t-test.

<u>Maximal voluntary capacity (MVC) Analysis</u>. We computed the participant’s initial maximal strength by averaging the maximal force produced during each MVC trial of *MVC 1*. We assessed participants’ strength throughout the experimental sessions (MVC_Post-1,_ MVC_Post-2_) by quantifying the maximal force produced in each MVC trial. We compared participants’ MVC_Baseline_ between biofeedback conditions using a paired t-test. We also assessed the effect of the *fatigue choice phase* on participants’ strength using generalized linear mixed-effects (GLME) analyses via the *fitglme* function in MATLAB. As recommended and to account for differences between participants^52^, we included a random intercept of each participant in all models. We tested whether the type of biofeedback reduced participants’ MVC (dependent variable) differently before (MVC_Post-1_) and after the *fatigue choice phase* (MVC_Post-2_).

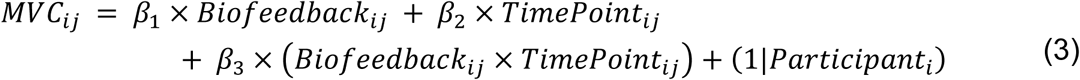

where *i* represents number of observations and *j* represents participants.

#### Subjective valuation of Effort Analysis

We used a two-parameter model to estimate participants’ subjective effort cost. We assumed a participant’s cost function *V(x)* for effort *x* as a power function of the form:

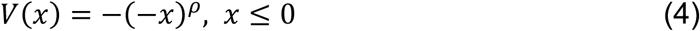

In this definition of effort cost, the effort level *x* is defined as negative, interpreted as force production being perceived as a loss. The parameter ρ represents sensitivity to changes in subjective effort value as the effort level changes. A large ρ represents a high sensitivity to increases in absolute effort level. ρ = 1 implies that subjective effort costs coincide with objective effort costs. Representing the effort levels as prospective costs, and assuming participants combine probabilities and utilities linearly, the relative value between the two effort options can be written as follows:

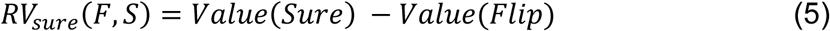

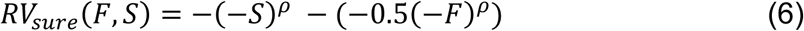

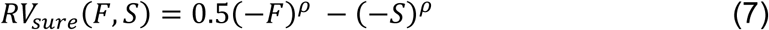

where RV_sure_ denotes the difference in value between the two options, and both F < 0 and S < 0 for all trials.

We then assume that the probability that a participant chooses the sure option for the *k*^th^ trial is given by the *softmax* function:

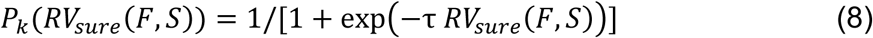

where τ is a non-negative temperature parameter representing the stochasticity of a participant’s choice (τ = 0 corresponds to random choice).

We used maximum likelihood to estimate parameters ρ and τ for each participant, based on 100 trials of effort choices (F, S), with a participant’s choice represented by y ∈ {0, 1}. Here, y = 1 indicates that the participant selected the sure option. This estimation was carried out by maximizing the likelihood function separately for each participant.

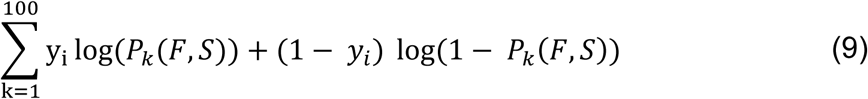

Parameter estimation was carried out separately for choices during the baseline and fatigue phases. This approach allowed us to obtain ρ_Baseline_ and ρ_Fatigue_ parameters for each participant in each biofeedback session.

We also used model-free metrics to examine whether there was a Biofeedback by Block interaction in the acceptance rate of the risky option by performing a GLME analysis.

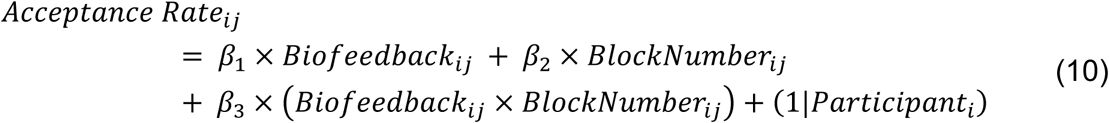

where *i* represents number of observations, *j* represents participants, and Block Number represents a numerical variable ranging from 0 to 9.

Lastly, to understand the effect of biofeedback condition on subjective effort cost over time, we performed a GLME analysis (Mixed-effect logistic regression) to examine whether the effect of biofeedback on subjective effort cost was influenced by trial number and differences in expected value (DEV).

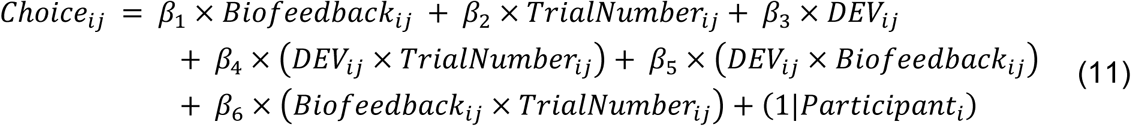

where *i* represents number of observations and *j* represents participants.

#### Fatigue Ratings Analysis

We used the numerical responses from each participant during the fatigue survey as our quantitative measure of fatigue ratings. To evaluate the level of fatigue experienced during each biofeedback condition, we conducted a GLME analysis. We examined whether the type of biofeedback provided affected the feeling of fatigue (dependent variable) across exertion blocks.

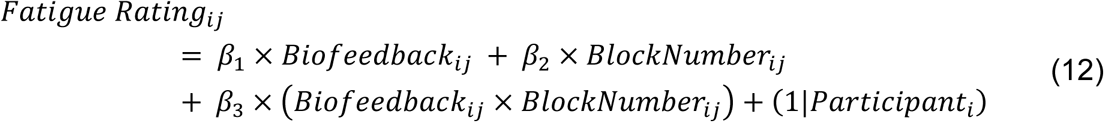

where *i* represents number of observations, *j* represents participants, and Block Number represents a numerical variable ranging from 0 to 9.

#### Assessment of Effort Analysis

We used the numerical responses from each participant during the effort survey as our quantitative measure of effort assessment. To quantify effort assessment during each biofeedback condition, we conducted a GLME analysis. We tested whether the type of biofeedback provided affected the effort assessment (dependent variable) across exertion blocks.

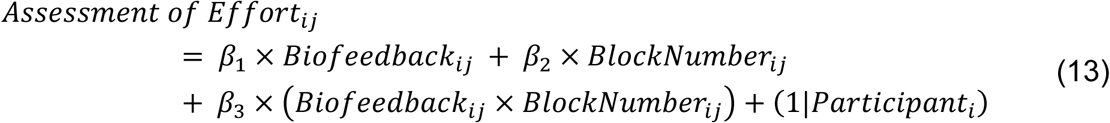

where *i* represents number of observations, *j* represents participants, and Block Number represents a numerical variable ranging from 0 to 9.

## Data availability

The data and analysis files will be permanently archived and publicly available for download on the Open Science Framework. We will provide the link upon acceptance of the manuscript for publication.

## Author contributions

ACM and VSC conceptualized the study. ACM developed the methodology under the supervision of VSC. ACM conducted the experiments. ACM, AK, and JLL analyzed the data under the supervision of VSC. VSC acquired the funding for the study. ACM wrote the original draft of the manuscript. ACM, AK, JLL, and VSC reviewed and edited the manuscript.

## Bibliography

1. Raizen, D. M. et al. Beyond the symptom: the biology of fatigue. SLEEP 46, zsad069 (2023).

2. Kluger, B. M., Krupp, L. B. & Enoka, R. M. Fatigue and fatigability in neurologic illnesses: Proposal for a unified taxonomy. Neurology 80, 409–416 (2013).

3. Penner, I.-K. & Paul, F. Fatigue as a symptom or comorbidity of neurological diseases. Nat. Rev. Neurol. 13, 662–675 (2017).

4. Hogan, P. S., Chen, S. X., Teh, W. W. & Chib, V. S. Neural mechanisms underlying the effects of physical fatigue on effort-based choice. Nat. Commun. 11, 4026 (2020).

5. Chong, T. T.-J. et al. Neurocomputational mechanisms underlying subjective valuation of effort costs. PLOS Biol. 15, e1002598 (2017).

6. Müller, T., Klein-Flügge, M. C., Manohar, S. G., Husain, M. & Apps, M. A. J. Neural and computational mechanisms of momentary fatigue and persistence in effort-based choice. Nat. Commun. 12, 4593 (2021).

7. Meyniel, F., Sergent, C., Rigoux, L., Daunizeau, J. & Pessiglione, M. Neurocomputational account of how the human brain decides when to have a break. Proc. Natl. Acad. Sci. 110, 2641–2646 (2013).

8. Hunter, S. K., Duchateau, J. & Enoka, R. M. Muscle Fatigue and the Mechanisms of Task Failure. Exerc. Sport Sci. Rev. 32, 44–49 (2004).

9. Enoka, R. M. & Duchateau, J. Muscle fatigue: what, why and how it influences muscle function. J. Physiol. 586, 11–23 (2008).

10. Bigland-Ritchie, B., Furbush, F. & Woods, J. J. Fatigue of intermittent submaximal voluntary contractions: central and peripheral factors. J. Appl. Physiol. 61, 421–429 (1986).

11. Enoka, M. & Stuart, G. Neurobiology of muscle fatigue.

12. Barry, B. K. & Enoka, R. M. The neurobiology of muscle fatigue: 15 years later. Integr. Comp. Biol. 47, 465–473 (2007).

13. Gandevia, S. C. Spinal and Supraspinal Factors in Human Muscle Fatigue. Physiol. Rev. 81, 1725–1789 (2001).

14. Klass, M., Lévénez, M., Enoka, R. M. & Duchateau, J. Spinal Mechanisms Contribute to Differences in the Time to Failure of Submaximal Fatiguing Contractions Performed With Different Loads. J. Neurophysiol. 99, 1096–1104 (2008).

15. Bigland-Ritchie, B. & Woods, J. J. Changes in muscle contractile properties and neural control during human muscular fatigue. Muscle Nerve 7, 691–699 (1984).

16. Stephan, K. E. et al. Allostatic Self-efficacy: A Metacognitive Theory of Dyshomeostasis-Induced Fatigue and Depression. Front. Hum. Neurosci. 10, (2016).

17. Casamento-Moran, A., Mooney, R. A., Chib, V. S. & Celnik, P. A. Cerebellar Excitability Regulates Physical Fatigue Perception. J. Neurosci. 43, 3094–3106 (2023).

18. Noakes, T. D. Fatigue is a Brain-Derived Emotion that Regulates the Exercise Behavior to Ensure the Protection of Whole Body Homeostasis. Front. Physiol. 3, (2012).

19. De Morree, H. M., Klein, C. & Marcora, S. M. Perception of effort reflects central motor command during movement execution. Psychophysiology 49, 1242–1253 (2012).

20. De Morree, H. M. & Marcora, S. M. Psychobiology of Perceived Effort During Physical Tasks. in Handbook of Biobehavioral Approaches to Self-Regulation (eds Gendolla, G. H. E., Tops, M. & Koole, S. L.) 255–270 (Springer New York, New York, NY, 2015). doi:10.1007/978-1-4939-1236-0_17.

21. Pageaux, B. Perception of effort in Exercise Science: Definition, measurement and perspectives. Eur. J. Sport Sci. 16, 885–894 (2016).

22. Hogan, P. S., Galaro, J. K. & Chib, V. S. Roles of Ventromedial Prefrontal Cortex and Anterior Cingulate in Subjective Valuation of Prospective Effort. Cereb. Cortex 29, 4277–4290 (2019).

23. Greenhouse-Tucknott, A. et al. Toward the unity of pathological and exertional fatigue: A predictive processing model. Cogn. Affect. Behav. Neurosci. 22, 215–228 (2022).

24. Sennesh, E. et al. Interoception as modeling, allostasis as control. Biol. Psychol. 167, 108242 (2022).

25. Sterling, P. Allostasis: A model of predictive regulation. Physiol. Behav. 106, 5–15 (2012).

26. Quigley, K. S., Kanoski, S., Grill, W. M., Barrett, L. F. & Tsakiris, M. Functions of Interoception: From Energy Regulation to Experience of the Self. Trends Neurosci. 44, 29–38 (2021).

27. Barrett, L. F. & Simmons, W. K. Interoceptive predictions in the brain. Nat. Rev. Neurosci. 16, 419–429 (2015).

28. Shaffer, C., Westlin, C., Quigley, K. S., Whitfield-Gabrieli, S. & Barrett, L. F. Allostasis, Action, and Affect in Depression: Insights from the Theory of Constructed Emotion. Annu. Rev. Clin. Psychol. 18, 553–580 (2022).

29. Barrett, L. F. The theory of constructed emotion: an active inference account of interoception and categorization. Soc. Cogn. Affect. Neurosci. nsw154 (2016) doi:10.1093/scan/nsw154.

30. Craig, A. D. How do you feel? Interoception: the sense of the physiological condition of the body. Nat. Rev. Neurosci. 3, 655–666 (2002).

31. (Bud) Craig, A., Interoception: the sense of the physiological condition of the body. Curr. Opin. Neurobiol. 13, 500–505 (2003).

32. Nethery, V. M. Competition between internal and external sources of information during exercise: influence on RPE and the impact of the exercise load. J. Sports Med. Phys. Fitness 42, 172–178 (2002).

33. Chen, M. J., Fan, X. & Moe, S. T. Criterion-related validity of the Borg ratings of perceived exertion scale in healthy individuals: a meta-analysis. J. Sports Sci. 20, 873–899 (2002).

34. Leocani, L., Colombo, B. & Comi, G. Physiopathology of fatigue in Multiple Sclerosis. Neurol. Sci. 29, 241–243 (2008).

35. Gonzalez Campo, C., et al. Fatigue in multiple sclerosis is associated with multimodal interoceptive abnormalities. Mult. Scler. J. 26, 1845–1853 (2020).

36. Krupp, L. Editorial. Mult. Scler. J. 12, 367–368 (2006).

37. Kuppuswamy, A., Clark, E. V., Turner, I. F., Rothwell, J. C. & Ward, N. S. Post-stroke fatigue: a deficit in corticomotor excitability? Brain 138, 136–148 (2015).

38. Hofman, M., Ryan, J. L., Figueroa-Moseley, C. D., Jean-Pierre, P. & Morrow, G. R. Cancer-Related Fatigue: The Scale of the Problem. The Oncologist 12, 4–10 (2007).

39. Morrow, G. R., Andrews, P. L., Hickok, J. T., Roscoe, J. A. & Matteson, S. Fatigue associated with cancer and its treatment. Support. Care Cancer 10, 389–398 (2002).

40. Dantzer, R., O’Connor, J. C., Freund, G. G., Johnson, R. W. & Kelley, K. W. From inflammation to sickness and depression: when the immune system subjugates the brain. Nat. Rev. Neurosci. 9, 46–56 (2008).

41. Townsend, L. et al. Persistent fatigue following SARS-CoV-2 infection is common and independent of severity of initial infection. PLOS ONE 15, e0240784 (2020).

42. Van Herck, M. et al. Severe Fatigue in Long COVID: Web-Based Quantitative Follow-up Study in Members of Online Long COVID Support Groups. J. Med. Internet Res. 23, e30274 (2021).

43. Al-Aly, Z., Xie, Y. & Bowe, B. High-dimensional characterization of post-acute sequelae of COVID-19. Nature 594, 259–264 (2021).

44. Gotaas, M. E., Stiles, T. C., Bjørngaard, J. H., Borchgrevink, P. C. & Fors, E. A. Cognitive Behavioral Therapy Improves Physical Function and Fatigue in Mild and Moderate Chronic Fatigue Syndrome: A Consecutive Randomized Controlled Trial of Standard and Short Interventions. Front. Psychiatry 12, 580924 (2021).

45. Fedorowski, A. et al. Cardiovascular autonomic dysfunction in post-COVID-19 syndrome: a major health-care burden. Nat. Rev. Cardiol. 21, 379–395 (2024).

46. Van Rhijn-Brouwer, F. C. C.-C. et al. Graded exercise therapy should not be recommended for patients with post-exertional malaise. Nat. Rev. Cardiol. 21, 430–431 (2024).

47. Castell, B. D., Kazantzis, N. & Moss-Morris, R. E. Cognitive behavioral therapy and graded exercise for chronic fatigue syndrome: A meta-analysis. Clin. Psychol. Sci. Pract. 18, 311–324 (2011).

48. Meyniel, F. et al. A specific role for serotonin in overcoming effort cost. eLife 5, e17282 (2016).

49. Thase, M. E., Fava, M., DeBattista, C., Arora, S. & Hughes, R. J. Modafinil Augmentation of SSRI Therapy in Patients with Major Depressive Disorder and Excessive Sleepiness and Fatigue: *A 12-Week, Open-label, Extension Study*. CNS Spectr. 11, 93–102 (2006).

50. Papakostas, G. I. et al. Resolution of Sleepiness and Fatigue in Major Depressive Disorder: A Comparison of Bupropion and the Selective Serotonin Reuptake Inhibitors. Biol. Psychiatry 60, 1350–1355 (2006).

51. Brainard, D. H. The Psychophysics Toolbox. Spat. Vis. 10, 433–436 (1997).

52. Seltman, H. J. Experimental Design and Analysis.

