## Supplementary Materials for "Neuromuscular Signals Shape Fatigue and Effort-Based Decision-Making in Humans"

**Affiliations:**

**This PDF file includes:**

Supplementary text and results
Supplementary Figures 1 to 2

### Supplementary Results

Baseline MVC. We compared participant's $MVC_{Baseline}$ between biofeedback conditions using a paired t-test to ensure that participants provided similar maximal strength in both sessions and found that $MVC_{Baseline}$ was similar between biofeedback sessions (Paired t-test: $t_{(1,23)} = -0.05$, $p = 0.96$).

Effect of the experimental manipulations on MVC. We found a significant main effect of Time Point (Linear mixed-effects model: $t_{(92)} = -1.98$, $p = 0.049$) indicating that the *fatigue choice phase* reduced participants' MVC (i.e., induced fatigability). However, we did not find a significant main effect of Biofeedback condition (Linear mixed-effects model: $t_{(92)} = -0.91$, $p = 0.36$), nor a Biofeedback by Time Point interaction (Linear mixed-effects model: $t_{(92)} = -0.54$, $p = 0.58$), indicating that both biofeedback conditions induced similar levels of fatigability.

Subjective valuation of Effort. First, we used a paired t-test to examine if subjective effort cost was similar between biofeedback conditions at baseline ($\rho_{Baseline}$) and found that $\rho_{Baseline}$ was not significantly different between biofeedback conditions (Paired t-test: $t_{(1,23)} = 0.38$, $p = 0.71$; **Supplementary Fig. 1**). Then, we used repeated measures ANOVA to examine how subjective effort cost (ρ) changed from the baseline to the fatigue choice phase and across biofeedback conditions. We found a significant main effect of Time (Repeated measures ANOVA: $F_{(1,23)} = 15.03$, $p < 0.001$) suggesting that ρ was significantly greater during the *fatigue choice phase* compared to baseline for both biofeedback conditions. In addition, we found a significant Biofeedback x Time interaction (Repeated measures ANOVA: $F_{(1,23)} = 6.41$, $p = 0.019$; **Fig. 4B**) demonstrating that force biofeedback resulted in a greater change in ρ than EMG biofeedback.

Acceptance rate of the risky option. We did not find a significant main effect of Biofeedback (Linear mixed-effects model: $t_{(434)} = -0.58$, $p = 0.57$), which indicated that at the first block, there were no significant differences in participants' choices across the two conditions. However, we found a significant main effect of Block Number (Linear mixed-effects model: $t_{(434)} = -4.27$, $p < 0.0001$), suggesting that the acceptance rate of the risky option (i.e., Flip) reduced across choice blocks during the *fatigue choice phase.* Importantly, we also found significant Biofeedback by Block interaction (Linear mixed-effects model: $t_{(434)} = 2.06$, $p = 0.04$; **Fig. 4C**) demonstrating that the acceptance rate of the risky option decreased the most during force biofeedback.

Subjective valuation of Effort as a function of Time and DEV. While we did not find a significant main effect of Biofeedback (Linear mixed-effects model: $t_{(4361)} = 0.11$, $p = 0.91$), we found a significant main effect of Trial Number (Linear mixed-effects model: $t_{(4361)} = -4.68$, $p < 0.0001$) suggesting that the acceptance of the risky option (i.e., Flip) reduced across choice trials during the *fatigue choice phase*, as well as a significant main effect of DEV (Linear mixed-effects model: $t_{(4361)} = 13.98$, $p < 0.0001$) suggesting that the acceptance of the risky option (i.e., Flip) increased with increasing DEV. We also found significant Biofeedback by Trial Number interaction demonstrating choice preferences changed the most across trials with force rather than EMG biofeedback (Linear mixed-effects model: $t_{(4361)} = 3.18$, $p = 0.001$, **Fig. 4D**).

Fatigue Ratings. We did not find a significant main effect of Biofeedback (Linear mixed-effects model: $t_{(1004)} = -1.58$, $p = 0.11$) nor a Biofeedback by Block interaction (Linear mixed-effects model: $t_{(1004)} = 0.64$, $p = 0.52$). However, we did find a significant main effect of Block Number (Linear mixed-effects model: $t_{(1004)} = 11.02$, $p < 0.001$) indicating that fatigue ratings increased with repeated effort exertion blocks. Together, these results suggest that both biofeedback conditions increased feelings of fatigue similarly throughout the *fatigue choice phase* (**Fig. 5A**).

Assessment of Effort. We did not find a significant main effect of Biofeedback (Linear mixed-effects model: $t_{(545)} = 0.52$, $p = 0.60$) nor Block Number (Linear mixed-effects model: $t_{(545)} = 0.67$, $p = 0.50$), neither a Biofeedback by Block Number interaction (Linear mixed-effects model: $t_{(545)} = -1.5$, $p = 0.13$) demonstrating that the *fatigue choice phase* did not heighten assessments of effort and that both biofeedback conditions resulted in similar levels of effort assessment (**Fig. 5B**).

### Supplementary Figures

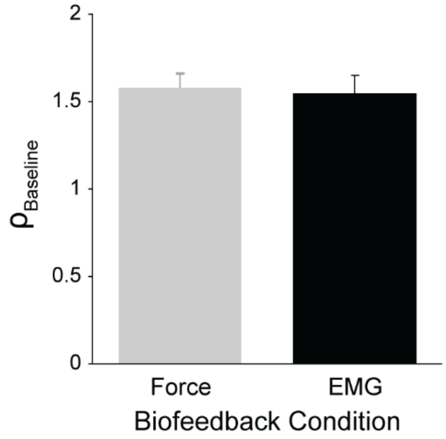


**Supplementary Figure 1: Subjective valuation of effort at baseline.** Mean subjective effort cost function at baseline ($\rho_{Baseline}$) for both biofeedback conditions, demonstrating that $\rho_{Baseline}$ was not significantly different between biofeedback conditions ($p$ = 0.71). Error bars indicate SEM.

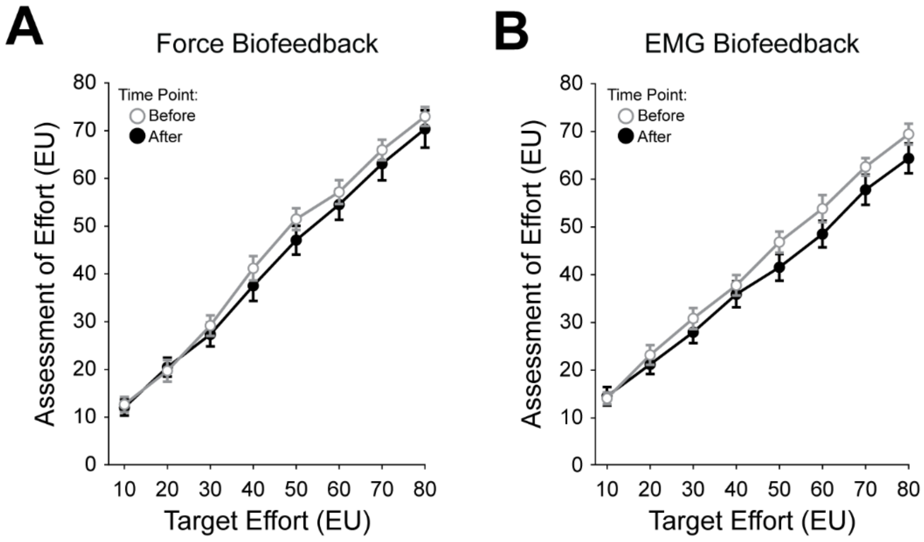


**Supplementary Figure 2: Assessment of effort before and after the fatigue choice phase. (A)** Mean assessment of effort as a function of target effort goal before (white) and after (black) the fatiguing paradigm with Force biofeedback. **(B)** Mean assessment of effort as a function of target effort goal before (white) and after (black) the fatiguing paradigm with EMG biofeedback. Error bars indicate SEM.
